# Molecular and physiological responses predict acclimation limits in juvenile brook trout (*Salvelinus fontinalis*)

**DOI:** 10.1101/2020.12.07.414821

**Authors:** Theresa E. Mackey, Caleb T. Hasler, Travis Durhack, Jennifer D. Jeffrey, Camille J. Macnaughton, Kimberly Ta, Eva C. Enders, Ken M. Jeffries

**Affiliations:** University of Winnipeg, Winnipeg, MB, Canada; University of Manitoba, Winnipeg, MB, Canada; Fisheries and Oceans Canada, Freshwater Institute, Winnipeg, MB, Canada

**Keywords:** transcriptomics, stress response, metabolic rate, temperature, ectotherm, freshwater fish

## Abstract

Brook trout (*Salvelinus fontinalis*) populations are at risk of exposure to high water temperatures in the species’ native range in eastern North America. We quantified the physiological and molecular responses of juvenile brook trout to six acclimation temperatures that span the thermal distribution of the species (5, 10, 15, 20, 23, and 25°C). Using quantitative PCR (qPCR), we measured the mRNA transcript abundance of temperature-induced cellular stress genes to identify a potential sub-lethal temperature threshold for brook trout between 20–23°C. Brook trout exhibited an upregulation of stress-related genes (*heat shock protein 90-beta*; *heat shock cognate 71 kDa protein*; *glutathione peroxidase 1*) and a downregulation of transcription factors and osmoregulation-related genes (*Na^+^/K^+^/2Cl^−^ co-transporter-1-a*; *nuclear protein 1*) at temperatures ≥20°C. We also used respirometry to assess the effects of the acclimation temperatures on oxygen consumption. Standard metabolic rate results indicated that energy expenditure was higher at temperatures ≥20°C. We then examined the effects of acclimation temperature on metabolic rate and blood plasma parameters in fish exposed to an acute exhaustive exercise and air exposure stress. Fish acclimated to temperatures ≥20°C exhibited elevated levels of plasma cortisol, muscle lactate, and plasma glucose after exposure to the acute stressors. After 24 h of recovery, fish showed longer metabolic recovery times at 15 and 20°C and cortisol levels remaining elevated at temperatures ≥20°C. Our findings suggest that brook trout may have a limited ability to acclimate to temperatures >20°C and increases in temperatures beyond 20°C may impact brook trout populations.

## Introduction

Climate change is altering thermal habitats of freshwater ectotherms, which can lead to a shift in species distributions, contribute to disease outbreaks, influence phenology, and decrease survival (Bassar et al., 2016; Hermoso, 2017; Krabbenhoft et al., 2017). It is estimated that 50% of freshwater species are threatened by climate change and associated warming temperatures (Darwall and Freyhof, 2016; Reid et al., 2019). For freshwater fishes, temperature is a ‘master’ abiotic factor and it affects most major physiological and ecological processes (Fry, 1947; Beitinger and Bennett, 2000; Somero, 2005). Increases in temperature that surpass the physiological limits of some fishes can lead to extirpation or extinction (Nogués-Bravo et al., 2018). Fishes differ greatly in their abilities to tolerate temperatures outside their thermal distribution (Rahel, 2002) and individual fish behaviourally thermoregulate to remain in waters with ‘preferred’ temperatures (Martins et al., 2011; Cott et al., 2015; Raby et al., 2018), and by selecting thermally heterogeneous microhabitats (Brett, 1971; Nevermann and Wurtsbough, 1994; Nielsen et al., 1994; Biro, 1998; Newell and Quinn, 2005). Ultimately, if a fish is unable to leave or acclimate to the temperature of the environment, exposure may lead to mortality.

The ability to acclimate to high water temperatures is an adaptive response that can mitigate against the effects of environmental change (Crozier and Hutchings, 2014). Acclimation is a reversible phenotypic change caused by exposure to a changing environmental condition (i.e., temperature) for a period of days to months (Hochachka and Somero, 2002; Havird et al., 2020). During acclimation, aquatic ectotherms undergo changes in biological processes, including those at a cellular level, to maintain homeostasis (Schreck and Tort, 2016). These changes at a cellular level can alter an aquatic ectotherm’s phenotype and potentially allow for an adaptive response to environmental change (i.e., altered expression of protein isoforms; Hochachka and Somero, 2002). Additionally, fluctuations in thermal acclimation ability demonstrate when sub-lethal thresholds begin to have adverse effects on fishes (Komoroske et al., 2015). Organisms that can acclimate to changing thermal conditions, are likely to contribute more to future generations, which could affect species at a population level (Schulte, 2014). Therefore, if aquatic ectotherms can successfully acclimate to warming temperatures, their populations may tolerate climate change-related temperature increases in their current distributions.

Brook trout (*Salvelinus fontinalis;* Mitchill, 1814) are a predominantly freshwater salmonid fish species in North America, with some populations facing the threat of extirpation caused by increases in water temperatures in their native range (Chadwick et al., 2015). Climate projections show that range losses for brook trout along their southern boundary could reach up to 49% by 2050 (Meisner, 1990a; Chu et al., 2005; Flebbe et al., 2006). Numerous studies have suggested that the preferred temperature (i.e., temperature at which fish congregate when placed within a thermal gradient in the lab) for brook trout is approximately 15°C (e.g., Graham, 1949; Fry, 1971; Cherry et al., 1977; Stitt et al. 2014), with an upper temperature limiting habitat use of 21–23.5°C in the wild (Meisner, 1990a; Benfey et al., 1997; DeWeber and Wagner, 2015; Chadwick et al., 2015; Chadwick and McCormick, 2017). Because of the risk of rising temperatures on some populations of brook trout, a better understanding of the acclimation limits of this species will aid in predicting the potential effects of future climate change on these populations.

Integrating whole-organism physiology with molecular techniques can help to identify the ability of a fish to tolerate or acclimate to elevated water temperatures. Whole-organism oxygen consumption levels, as an estimate of metabolic rate, increase with temperature, suggesting elevated energetic demands being placed on the organism (Pörtner, 2001, 2002; Pörtner et al., 2017). Additionally, when fish are exposed to a thermal disturbance or stressor, as part of the generalized stress response, cortisol, the main glucocorticoid in fish, is released as an end product of the hypothalamic-pituitary-inter-renal (HPI) axis (Wendelaar Bonga, 1997; Barton, 2002). Increases in circulating levels of cortisol result in the increased mobilization of energy resources, such as glucose (Wendelaar Bonga, 1997; Barton, 2002). In response to an increase in temperature, cellular-level responses also occur and include changes in the expression of genes responsible for various processes that are necessary to help the fish react to the stressor. Therefore, transcriptomics can be useful in determining thermal tolerance thresholds of individuals (Connon et al., 2018) and has contributed to advances in understanding the cellular processes behind whole-organism physiological responses (Miller et al., 2014; Evans, 2015). For example, proteins such as heat shock proteins, are useful to measure as an indicator of a response to a temperature stressor and to monitor acute and chronic changes in the organism. There can be isoforms of some heat shock proteins that may be continuously expressed over time (constitutive) and useful to indicate the thermal acclimation response, in addition to isoforms that can be induced as a response to an acute stressor (inducible) where transient increases occur during an acute response (Iwama et al., 1998). Therefore, changes in the expression of some genes, as estimated by the abundance of mRNA transcripts, also provides information about the acute and chronic effects of temperature at the cellular level. Some cellular responses may be altered or peak prior to detrimental physiological or whole organism changes (Jeffries et al., 2014; Jeffries et al., 2018), which can help identify sub-lethal thresholds beyond which can lead to a negative impact on the individual (Connon et al., 2018).

In this study, we examined the effects of temperature acclimation on the cellular and physiological processes in juvenile brook trout to identify sub-lethal thresholds and acclimation limits for this species. We also examined the effects of acclimation temperature on blood stress indices and metabolic rate following exposure to acute exhaustive exercise and air exposure stressors, as well as metabolic rate recovery following the stressor. We tested the hypothesis that environmental temperature acclimation limits of a species can be predicted by identifying sub-lethal thresholds at the transcriptome level and the ability to recover from acute stress events. Previous studies have suggested that temperatures between 20–23°C lead to reduced physiological performance in brook trout (Smith and Ridgway, 2019; Morrison et al., 2020). Brook trout in the present study were acclimated for 21 days to one of six different temperatures (5, 10, 15, 20, 23, and 25°C) that span the thermal distribution of the species. We predicted that mRNA abundance of transcripts associated with the cellular stress response (i.e., reducing damage to cellular proteins due to heat stress, preventing damage from reactive oxygen species, and regulating cell growth) would be elevated when fish are acclimated to temperatures beyond a sub-lethal threshold (i.e., >20°C in brook trout). Additionally, we predicted that an exhaustive exercise and air exposure treatment would increase activation of the stress response (i.e., plasma cortisol, plasma glucose, plasma osmolality, and muscle lactate) at temperatures ≥20°C. Further, we predicted that metabolic rate would increase with temperature, where those fish exposed to the higher temperature groups (≥20°C) would experience the longest metabolic recovery time due to increased energetic demands. In this study, we showed that brook trout have a reduced ability to acclimate to temperatures ≥20°C as supported by changes in mRNA abundance, standard metabolic rate, and their ability to recover to exercise stress.

## Materials and methods

### Study animals

The juvenile brook trout (mass = 38.3 ± 1.7 g, fork length = 14.5 ± 0.2 cm) used in this study were first generation (F1) brook trout originally obtained from the Whiteshell Fish Hatchery in eastern Manitoba, Canada. In 2016, brood stock brook trout were bred at the Fisheries and Oceans Canada (DFO) Freshwater Institute in Winnipeg, Manitoba, Canada. After hatching (January 2017) and when past the swim-up stage, fish were moved to one of two aerated 600 l circular flow-through tanks at approximately 10°C. Fish were fed *ad libitum* with commercial pellet fish food (EWOS Pacific: Complete Fish Feed for Salmonids, Cargill, Winnipeg, Manitoba, Canada) for a 35-week rearing period. All methods were approved by the Freshwater Institute Animal Care Committee (FWI-ACC-AUP-2018-02/2019-02).

### Temperature treatment

Juvenile brook trout (*n* = 140) were haphazardly netted from the general population tank and placed into 200 l aerated, flow-through tanks and exposed to one of six temperatures (5, 10, 15, 20, 23, and 25°C; *n* = 50 per temperature tank) for 21 days. Due to logistic constraints, temperature exposures were staggered across four months in 2018–19: 10°C beginning on October 11, 25°C on October 23, 23°C on November 2, 20°C on November 16, 15°C on December 17, and 5 °C on January 1. On the first day of each temperature treatment, fish were transferred to a 200 l acclimation tank at 10–11°C and were given 1 day to recover from the handling stress. The water temperature was then gradually adjusted to the assigned treatment temperature at a rate of 1.5–2°C day^−1^ using heating or cooling coils that were placed in an auxiliary tank plumbed to the holding tank. Once the treatment temperature was reached, fish remained at the temperature for a 21-day acclimation period (Beitinger et al., 2000). Throughout the treatment period, the water temperature of the holding tank was measured using a HOBO Tidbit v2 Sensor (ONSET Computer Corporation, Bourne, Massachusetts, USA) and controlled with WitroxCTRL software (Loligo^®^ Systems, Tjele Denmark), where it fluctuated daily by ± 1.5°C of the treatment temperature to simulate diurnal temperature changes. A 12:12 hour day-night cycle was used throughout the experiment (65 min of dawn and dusk, full-light starting at 07:05, and full dark at 19:05). Dissolved oxygen was kept above 7 mg l^−1^ throughout the experiment. After the 21-day acclimation period, fish were designated to one of three groups: “unhandled, acute-stress, or acute-recovery”. Those termed the “unhandled” group were immediately euthanized and sampled (see below; *n* = 10). Those fish in the “acute” groups were exposed to a chase test and air exposure and either euthanized and sampled 30 min after the stressor exposure (termed the “acute-stress” group; *n* = 8) or placed in a respirometer for 24 h and euthanized afterwards (termed the “acute-recovery” group; *n* = 8). Fish from the 25°C group were not exposed to the full 21-day treatment period, as fish exhibited potential fungal infections, reduced feeding, and mortality. Therefore, the 25°C treatment group was sampled for tissues after 11 days (see below) and were not subjected to the acute stress experiments.

### Tissue sampling of unhandled group

Fish in the unhandled group (*n* = 60; *n* = 10 per treatment group) were individually euthanized in a buffered tricaine methanesulfonate solution [MS-222] (300mg l^−1^; buffered with 600 mg l^−1^ of sodium bicarbonate NaHCO_3_) with water at the same temperature as their treatment. Fish were measured for length and body mass prior to tissue sampling. Blood was collected by severing of the caudal fin and using ammonium-heparinized capillary tubes (Fisherbrand^®^, Fisher Scientific, Pittsburgh, Pennsylvania, USA). Whole-blood glucose was immediately measured using a UltraMini^®^ Glucose Meter (OneTouch^®^, LifeScan Canada, Burnaby, British Columbia, Canada) after which blood samples were centrifuged at 3000 × *g* for 6 min. Plasma was removed, flash frozen in liquid nitrogen, and stored at −80°C until analysis. The second two gill arches from the left side of each fish and the liver were sampled and placed in RNA*later*™ (Invitrogen™, Carlsbad, California, USA) and stored at 4°C overnight prior to storage at −80°C. White muscle was taken from the fish’s right side and placed in liquid nitrogen and stored at −80°C for measurement of muscle lactate.

### Acute stress and recovery

For the “acute-stress” and “acute-recovery” groups, another subset of fish from each acclimation temperature, with the exception of the 25°C group (see above), underwent an acute 2 min chase (e.g., Suski et al., 2006) and 5 min air exposure (e.g., Gingerich et al., 2007) and/or a recovery trial (*n* = 80; *n* = 8 per temperature and acute stress group group). The chase test consisted of placing individual fish into a bucket (diameter ≈ 17 in) with 15–20 l of water at the acclimation temperature. There was a pipe in the centre of the bucket to force the fish to swim around the perimeter of the container. Once the water was added to the bucket, the bucket was plugged, and the fish was manually chased using a small net. After 2 min, fish were netted out of the water and air exposed for 5 min. During the air exposure, fish were measured for length and mass. Following the acute stressor exposure, eight fish were placed into a holding tank at their acclimation temperature for 30 min to allow plasma cortisol and glucose to reach the putative peak values (Biron and Benfey, 1994; Benfey and Biron, 2000) before being euthanized and sampled for blood and white muscle as described above for unhandled individuals.

The remaining eight fish were placed in an intermittent-flow respirometry system, in a water bath held at the same temperature as the acclimation treatment, for 24 h to record oxygen consumption. Intermittent flow respirometry was used to quantify oxygen consumption as an estimate of metabolic rate. Oxygen consumption was measured by in-line probes connected to respirometry chambers (Presence, Regensburg, Germany) and automatically calculated by AutoResp software (Loligo Systems, Viborg, Denmark). To validate the quality of measurements, r^2^ values for rates of oxygen decline were also automatically generated. Only r^2^ values above 0.9 were used for final analysis of standard metabolic rate (SMR) and maximum metabolic rate (MMR). For SMR calculations, the lowest 20^th^ quantile of oxygen consumption rate (*M*O_2_ estimates were used after removing the first 10 h of measurements to ensure fish were at minimum oxygen consumption levels (Norin and Clark, 2016). Following this, the ‘FishMO2’ package in R (Chabot et al., 2016) was used to analyze *M*O_2_ estimates over time including the calculation of SMR and plotting of r^2^ values for each individual at each temperature to include those with a r^2^ > 0.9. Background respiration from microbial respiration (BOD) was also estimated by including an empty respirometry chamber in each trial and oxygen consumption values from these chambers were subtracted from the SMR measurements. MMR was estimated using a protocol of a 2 min chase test and 5 min air exposure. Three measurements of oxygen consumption were taken to estimate MMR when each fish was first placed into the respirometry chamber post exercise and an air exposure event (Norin and Clark, 2016). Aerobic scope (AS) was calculated by subtracting SMR estimates from MMR for each fish, and recovery time was determined as the time from when the fish was placed into the respirometer to the time until metabolic rate first began to stabilize (Cooke et al., 2014).

### Physiological assays

Blood plasma samples were used to measure cortisol, glucose, and osmolality. Plasma cortisol levels were quantified using an enzyme-linked immunosorbent assay (ELISA; 1:50 dilution; Neogen Corporation, Lexington, Kentucky, USA), previously validated for use in other salmonids (e.g., Jeffries et al., 2012b; Sopinka et al., 2016). The plasma osmolality was determined using a VAPRO vapour pressure osmometer (Wescor Inc., Logan, Utah, USA). Plasma glucose was quantified using a hexokinase kinetic glucose assay that was adapted for a 96-well plate (Treberg et al., 2007) where plasma samples were 1:29. Results from the hexokinase glucose assay were used to develop a correction factor for the whole-blood glucose levels measured using the UltraMini^®^ Glucose Meter (see above). Glucose values from the hexokinase kinetic assay and the glucose meter were compared using a linear regression and the resulting equation of the line was used to correct whole-blood values measured with the handheld meter (Fig. S1).

The white muscle was used to determine the amount of muscle lactate. White muscle samples were first powdered in liquid nitrogen using a mortar and pestle and tissue metabolites were extracted using an 8% perchloric acid solution mixed with ethylenediaminetetraacetic acid (EDTA), which was later neutralized using a base solution (mixture of sodium hydroxide, sodium chloride, and imidazole) to a pH between 7 and 8 (Booth et al. 1995). After metabolite extraction, lactate concentrations were determined using an enzymatic assay that utilized the reaction of converting lactate to pyruvate using NAD^+^ (nicotinamide adenine dinucleotide) and lactate dehydrogenase (Lowry and Passonneau, 1972; Gutman and Wahlefeld, 1974).

### Quantitative PCR

Total RNA was extracted from the gill and liver tissues using a Qiagen RNeasy Plus Mini Kit (Qiagen, Toronto, ON, CA) following manufacturer’s protocols. The RNA samples were checked for purity (A260/A280, A260/A230) and concentration using a NanoDrop One Spectrophotometer (ThermoFisher Scientific, Wilmington, DE, USA). The integrity of the RNA was assessed by electrophoresis on a 1% agarose gel. One μg of total RNA was reverse transcribed into cDNA using the Qiagen QuantiTect Reverse Transcription Kit (Qiagen, Valencia, California, USA) following manufacturer’s protocols, with the exception that the total volume was scaled to 32 μl.

All forward and reverse quantitative PCR (qPCR) primers (Table 1) were designed using Primer Express 3.0.1 (Applied Biosystems, ThermoFisher Scientific, Wilmington, DE, USA). Primers were designed using sequences from the brook trout transcriptome from Sutherland et al. (2019) (Table S1). Primers were designed for 13 target genes: *Na^+^/K^+^-transporting ATPase subunit alpha-3* (*atp1a3*), *cystic fibrosis transmembrane conductance regulator* (*cftr*), *cold-inducible RNA-binding protein* (*cirbp*), *glucose-6-phosphatase* (*g6pc*), *glutathione peroxidase-like peroxiredoxin* (*gpx1*), *ATP-sensitive inward rectifier K^+^ channel 8* (*irk8*), *heat shock cognate 71 kDA protein* (*hspa8*), *heat shock protein 90-beta-1* (*hsp90ab1*), *Na^+^/K^+^/2Cl^−^ co-transporter-1-a* (*nkcc1a*), *nuclear protein 1* (*nupr1*), *serpin h1* (*serpinh1*), and *B and E 1 subunits of V-type H-ATPase* (*vatb* and *vate1*) (Table 1). Primers were designed for three reference genes, *60s ribosomal protein L7* and *L8* (*rpl7* and *rpl8*) and *40s ribosomal protein S9* (*rps9*) (Table 1). Primer efficiencies were tested by generating standard curves using cDNA synthesized from the RNA pooled from 6 individuals from the treatment groups. Each 12 μL qPCR reaction consisted of 1 μL of a 1:10 dilution of cDNA, 500 nM forward and reverse primer, 6 μL of PowerUP SYBR Green Master Mix (Applied Biosystems, Thermo Fisher Scientific, Wilmington, DE, USA) and 4.8 μL RNase-free water. The qPCR reactions were run on a QuantStudio 5 Real-Time PCR System (Thermo Fisher Scientific, Life Technologies Corporation, Carlsbad, California USA) in 384 well plates. Target mRNA levels were normalized to the three reference genes using the 2^−ΔCt^ method (Livak and Schmittgen, 2001). The stability of the reference genes across treatments were confirmed using a pair-wise comparison with BestKeeper Version 1 (Pfaffl et al., 2004).

**Table 1.**
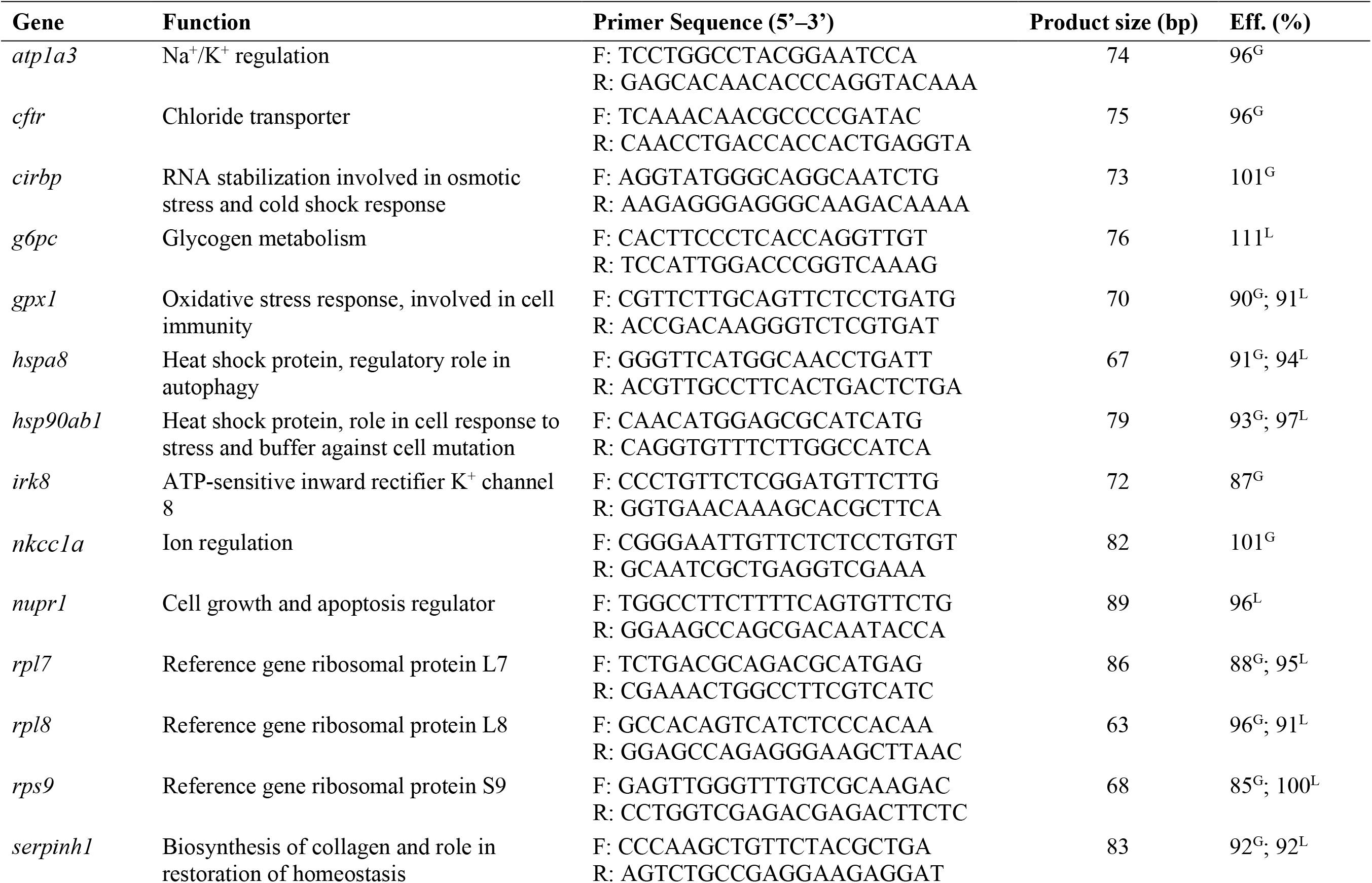
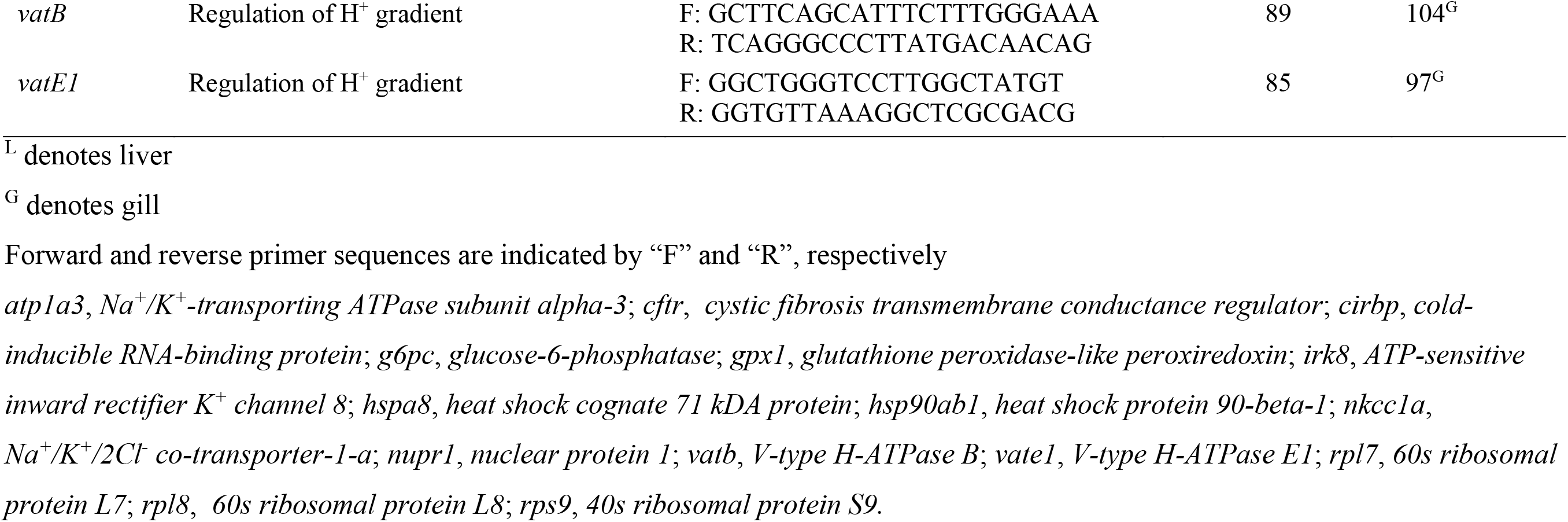
Primer sequences for quantitative PCR in brook trout (*Salvelinus fontinalis*).

### Statistical analysis

To determine whether mass was a significant factor contributing to physiological response variables (plasma cortisol, plasma glucose, tissue lactate, osmolality), a two-way analysis of covariance (ANCOVA) was initially run, but mass was shown to have no effect. For metabolic measurements (SMR and MMR), mass is used in the calculation of these metrics and therefore mass specific effects are not a factor. Therefore, the effect of acclimation temperature and treatment (unhandled, acute stress, acute recovery) on the physiological response variables (plasma cortisol, plasma glucose, tissue lactate, osmolality) were examined using two-way analyses of variance (ANOVAs) followed by Tukey’s honestly significant difference (HSD) post-hoc tests. Assumptions of normality and equal variance were assessed using a Shapiro-Wilk normality test and a Levene’s test, respectively. If assumptions of normality and equal variance were not met, a generalized linear mixed effects model (glmm; R Core Team, 2019) was used. In the above analyses our focus included the effects of the exhaustive exercise and air exposure stressor within an acclimation temperature, this was also the focus of the post-hoc tests (TukeyHSD or glmm). Therefore, the effects of the stressor within an acclimation temperature were the interactions we reported.

To determine the effect of acclimation temperature on the mRNA abundance and oxygen consumption parameters (SMR, MMR, time to recovery, and AS), ANOVAs were used followed by Tukey’s HSD post-hoc tests. If data failed to meet the assumptions of the ANOVA (see above), a Kruskal-Wallis test was run, followed by a Dunn’s post-hoc test.

All statistical analyses were run in R v.1.2.5033 (R Core Team, 2019). The level of significance (α) was 0.05 when one variable was analyzed (i.e., mRNA abundance). For multiple comparison tests (i.e., treatment and temperature), a Bonferroni corrected level of significance was used.

## Results

### Chronic temperature exposure effects on mRNA abundance

Gill mRNA abundance differed across acclimation temperatures for genes associated with thermal and oxidative stress (Fig. 1). The abundance of *nkcc1a* mRNA in the gill of fish held at 23 and 25°C was three-fold lower than at 5 and 10 °C (Fig. 1C; one-way ANOVA; *F_(5,59)_ =*5.069; *P* < 0.001). The abundance of *gpx1* mRNA was two-fold higher in the gill of fish held at 23 and 25°C compared with temperatures at and below 15°C (Fig. 1A; one-way ANOVA; *F_(5,59)_* = 5.735; *P* < 0.001). Conversely, the abundance of *hsp90ab1* mRNA was not significantly elevated in fish gill until the acclimation temperature reached 25°C, with a two-fold increase in comparison to the three coldest temperature treatments (Fig. 1B; one-way ANOVA; *F_(5,59)_* = 4.321; *P* = 0.002).

**Figure 1.**
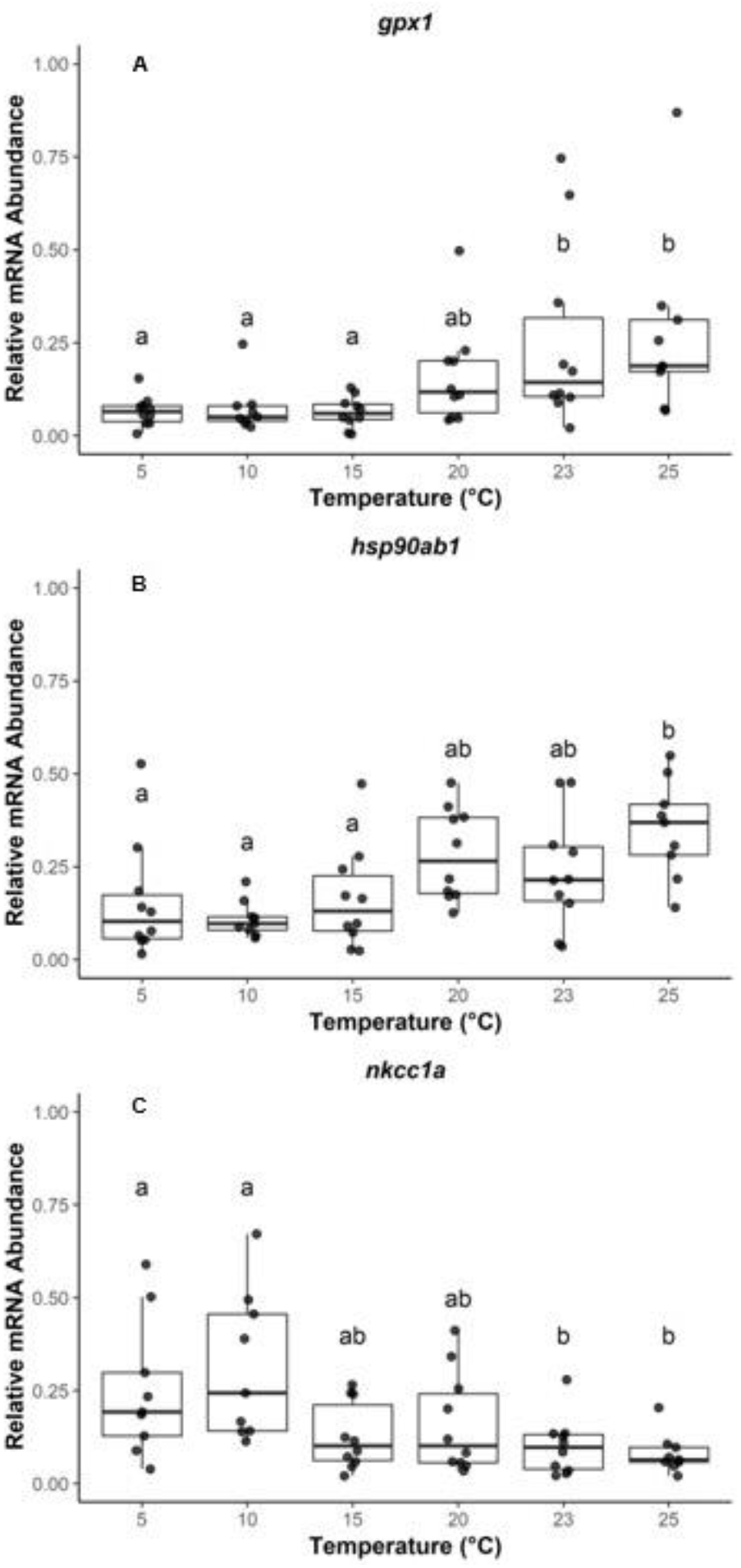
Transcript abundance of thermal stress biomarkers in gill tissue for juvenile brook trout (*Salvelinus fontinalis*) held at temperatures spanning their thermal distribution (*n* = 8–10). Fish were held for 21 days at the respective acclimation temperature, with the exception of fish held at 25°C, where fish were sampled after 11 days (see text for details). Groups that do not share a letter are significantly different from one another (one-way ANOVA, *P* < 0.05; see Table S2). Horizontal bars in the boxplot represent the median response value and the 75 and 25% quartiles. Whiskers represent ± 1.5 times the interquartile range, and each dot represents and individual response value. *gpx1*, *glutathione peroxidase-like peroxiredoxin*; *hsp90ab1*, *heat shock protein 90-beta-1*; *nkcc1a*, *Na^+^/K^+^/2Cl^−^ co-transporter-1-a*.

In liver tissue, the mRNA abundance of fives genes displayed significant responses (Fig. 2). The mRNA abundance of heat shock proteins *hspa8* (*hsc70*) and *hsp90ab1* was significantly elevated in the liver of fish held at 23 and 25°C, with a 4-fold (Fig. 2C; one-way ANOVA; *F_(5,59)_* = 8.828; *P* < 0.001) and 6-fold increase (Fig. 2D; one-way ANOVA; *F_(5,59)_* = 15.14; *P* < 0.001), respectively, compared to temperatures at and below 15°C. The mRNA abundance of *gpx1* was elevated by 4-fold in the liver of fish held at 5°C compared to those held at and above 20°C (Fig. 2B; one-way ANOVA; *F_(5,59)_* = 4.924; *P* = 0.001). The mRNA abundance of *g6pc* was highest in the liver of fish held at 20°C and 3-fold higher than those held at 5°C (Fig. 2A; one-way ANOVA; *F_(5,59)_* = 3.123; *P* = 0.016). Lastly, *nupr1* mRNA abundance was significantly elevated by 2-fold in fish held at 20°C compared to 25°C (Fig. 2E; one-way ANOVA; *F_(5,59)_* = 4.918; *P* = 0.001).

**Figure 2.**
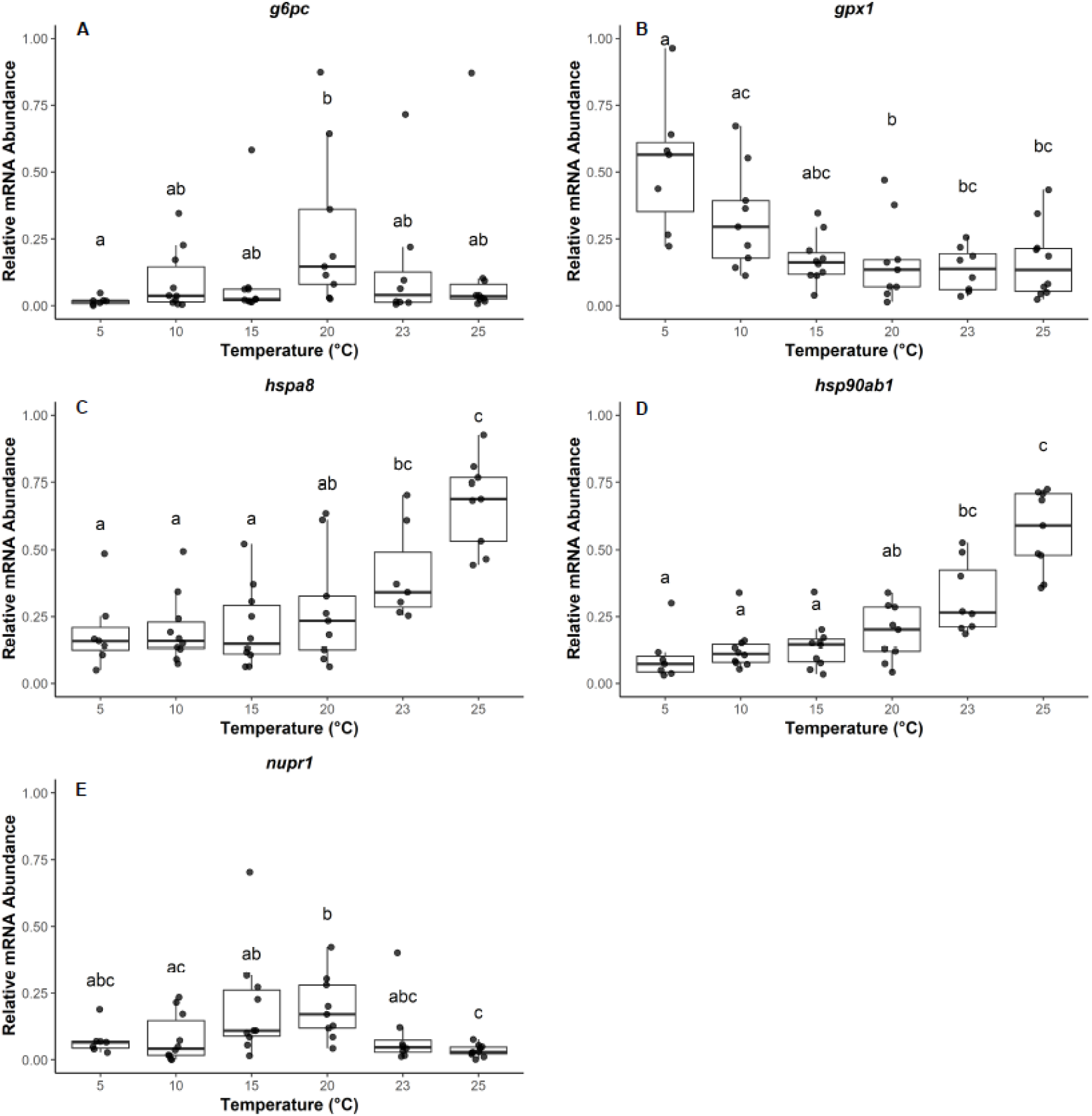
Transcript abundance of thermal stress biomarkers in liver tissue for juvenile brook trout (*Salvelinus fontinalis*) held at temperatures spanning their thermal distribution (*n* = 8–10). Fish were held for 21 days at the respective acclimation temperature, with the exception of fish held at 25°C, where fish were sampled after 11 days (see text for details). Groups that do not share a letter are significantly different from one another (one-way ANOVA, *P* < 0.05; see Table S2). Horizontal bars in the boxplot represent the median response value and the 75 and 25% quartiles. Whiskers represent ± 1.5 times the interquartile range, and each dot represents and individual response value. *g6pc*, *glucose-6-phosphatase*; *gpx1*, *glutathione peroxidase-like peroxiredoxin*; *hspa8*, *heat shock cognate 71 kDA protein*; *hsp90ab1*, *heat shock protein 90-beta-1*; *nupr1*, *nuclear protein 1*.

### Chronic temperature exposure effects on the acute stress response

Chasing followed by an air exposure had a significant effect on physiological variables associated with the stress response in juvenile brook trout. Regardless of acclimation temperature, muscle lactate was elevated 30 min following stressor exposure by an average of 1.8 times compared to fish sampled directly out of the acclimation tanks (i.e., unhandled) (Fig. 3A; two-way ANOVA, Treatment × Temperature; *F_(8)_* = 0.513; *P* < 0.001). Muscle lactate levels significantly returned to or below (for 23°C) pre-stressed levels following 24 h of recovery across all acclimation temperatures, except at 5°C, where lactate levels were not significantly different from pre- or post-stress levels. Fish held at 23°C exhibited the lowest muscle lactate level after 24 h of recovery from the stressor (6.2 μmol g^−1^ ± 0.8).

**Figure 3.**
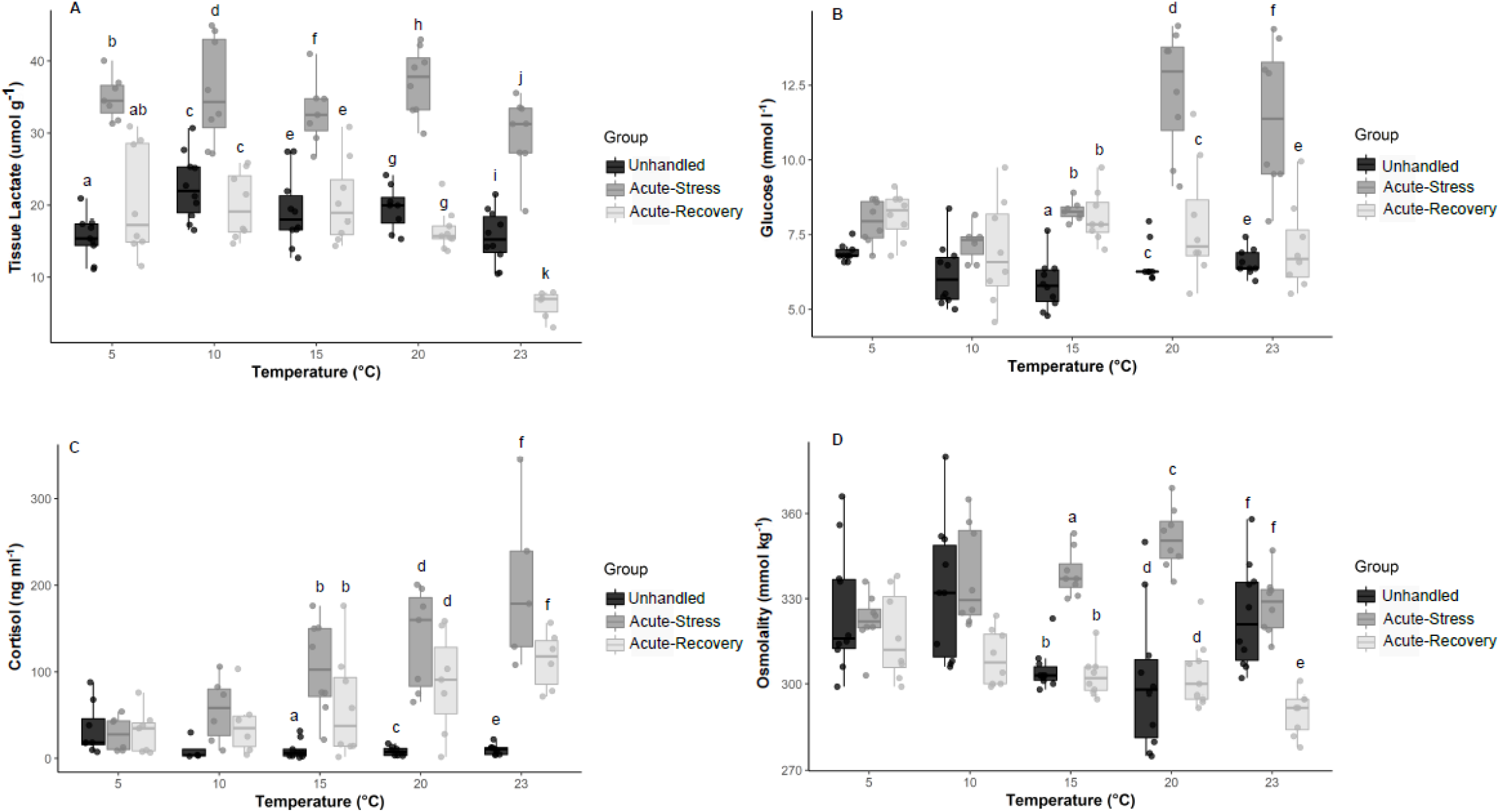
Physiological parameters collected from juvenile brook trout (*Salvelinus fontinalis*) held at temperatures spanning their thermal distribution for 21 days. Muscle lactate (A), plasma glucose (B), plasma cortisol (C), and plasma osmolality (D) were measured in fish directly sampled from the acclimation tank (unhandled; *n* = 10), 30 min (acute-stress, *n* = 8) and 24 h (acute-recovery; *n* = 8) after exposure to an acute stressor, consisting of 3 min of chasing and 5 min of air exposure. Within an acclimation temperature, groups that do not share a letter are significantly different from one another. For muscle lactate, plasma glucose and plasma cortisol, data were analyzed using a two-way ANOVA (*P* < 0.05; see Table S3). Plasma osmolality was analyzed using a glmm (*P* < 0.05; see Table 2). Horizontal bars in the boxplot represent the median response value and the 75 and 25% quartiles. Whiskers represent ± 1.5 times the interquartile range, and each dot represents and individual response value.

Acclimation temperatures also had a significant effect on both the plasma cortisol and glucose response to an acute stressor. For fish exposed to colder temperatures (5 and 10°C), exposure to the acute stressor had no significant effect on either plasma cortisol (Fig. 3C) or glucose (Fig. 3B) levels 30 min post-stressor exposure or following 24 h of recovery. However, fish exposed to warmer temperatures (15, 20, and 23°C) exhibited significantly elevated plasma cortisol and glucose levels following stressor exposure. Plasma cortisol levels remained elevated 24 h post-stressor exposure in these fish exposed to warmer temperatures (two-way ANOVA, Treatment × Temperature; *F_(8)_* = 4.31; *P* = 0.001). Plasma glucose levels returned to pre-stressed levels (i.e., unhandled) following 24 h of recovery for fish held at 20 and 23°C, but not for those held at 15°C (two-way ANOVA, Treatment × Temperature; *F_(8)_* = 6.929; *P* < 0.001).

Similarly, plasma osmolality did not differ significantly among groups (i.e., unhandled, acute-stress, acute-recovery) for fish held at colder acclimation temperatures (5 and 10°C; Fig. 3D; Table 2). Plasma osmolality increased by approximately 1.1 and 1.2-fold in response to stressor exposure for fish exposed to 15°C and 20°C, respectively (Table 2). At 23°C, fish had significantly lower plasma osmolality 24 h post-stressor exposure, but levels did not differ significantly between the unhandled fish and fish exposed to the acute stressor (Table 2).

**Table 2.**
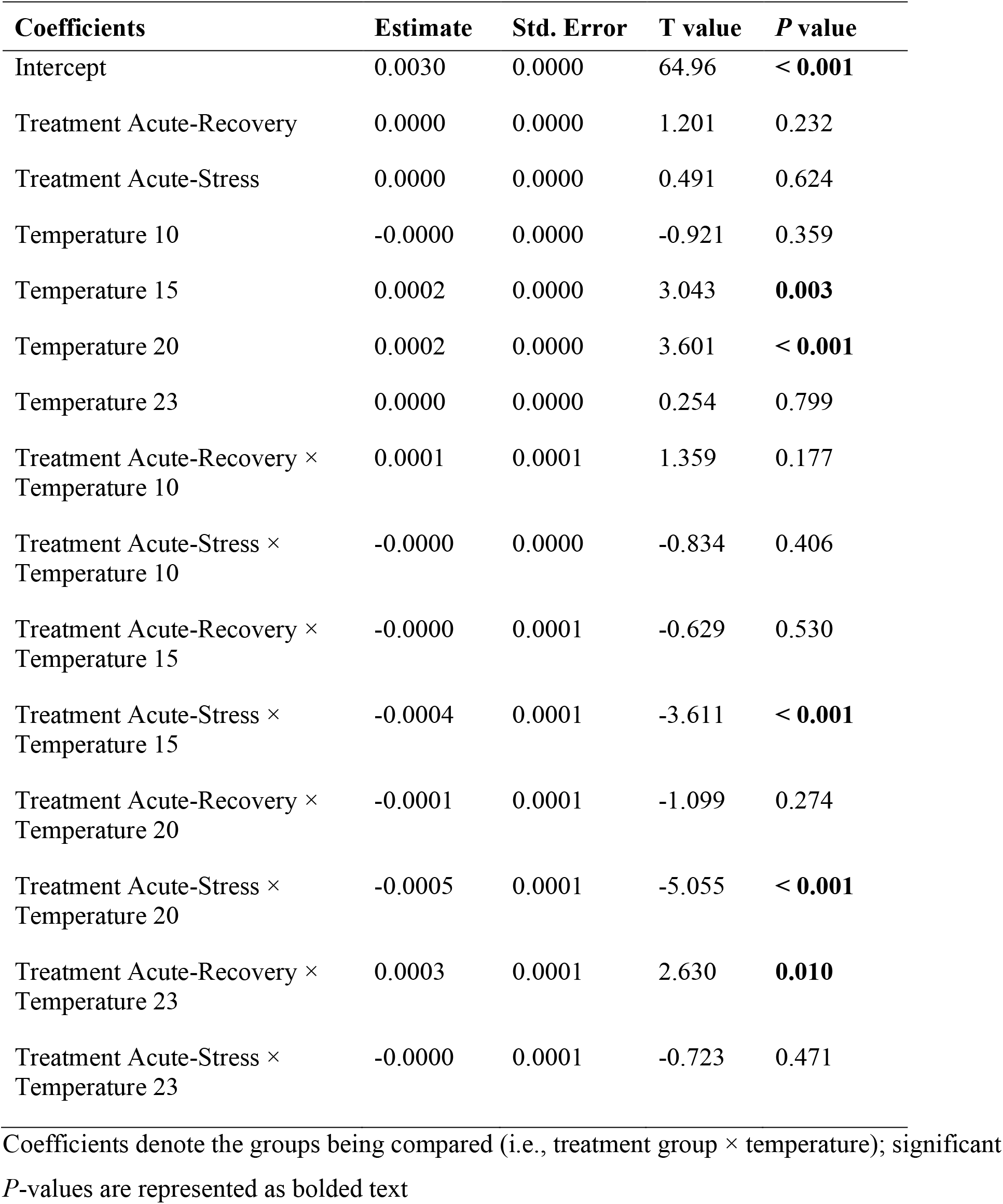
Results of the generalized linear model for plasma osmolality of juvenile brook trout (*Salvelinus fontinalis*) acclimated to five temperatures (5, 10, 15, 20, and 23°C) and exposed to one of three treatments (unhandled, acute stress, and acute-stress recovery).

### Chronic temperature exposure effects on metabolic rate

Standard metabolic rate of juvenile brook trout was unaffected until acclimation temperatures reached 20°C (Fig. 4A; Kruskal-Wallis; χ^2^ _(*4,40*)_ = 30.43; *P* < 0.001). Fish SMR was further increased by approximately 1.4-times in fish held at 23°C compared to 20°C. Maximum metabolic rate was highest in juvenile brook trout held at 23°C and lowest for fish held at 5°C (Fig. 4B; one-way ANOVA; *F_(4,40)_* = 8.84; *P* < 0.001). Aerobic scope was not significantly affected by acclimation temperature (Fig. 4D; one-way ANOVA; *F_(4,40)_* = 1.29; *P* = 0.292). Recovery time was approximately twice as long for fish exposed to 5, 15, and 20°C compared to 10 and 23°C (Fig. 4C; one-way ANOVA; *F_(4,40)_* = 27.69; *P* < 0.001).

**Figure 4.**
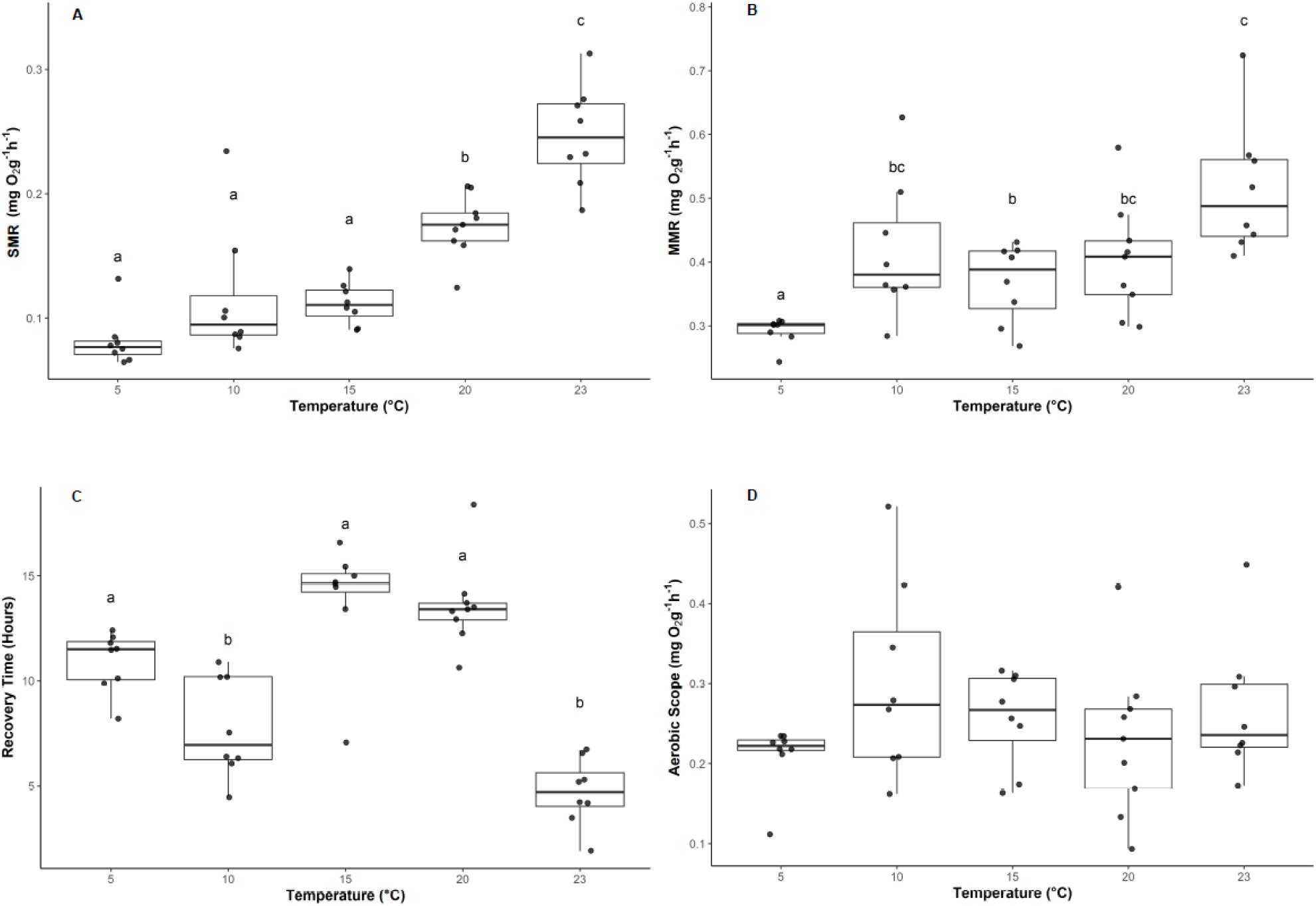
Metabolic and recovery parameters collected from juvenile brook trout (*Salvelinus fontinalis*) held at temperatures spanning their thermal distribution (*n* = 8). Standard metabolic rate (SMR; A), maximum metabolic rate (MMR; B), recovery time in hours post stress events (C), and aerobic scope (AS; D). Groups that do not share a letter are significantly different from one another (Kruskal-Wallis [SMR only], one-way ANOVA, *P* < 0.05; see Table S3). Horizontal bars in the boxplot represent the median response value and the 75 and 25% quartiles. Whiskers represent ± 1.5 times the interquartile range, and each dot represents and individual response value.

## Discussion

Our study assessed the ability of juvenile brook trout to acclimate to temperatures that span the thermal distribution of the species. It was evident that the physiology of juvenile brook trout was impacted at acclimation temperatures ≥20°C, as indicated by the mRNA transcript levels of genes associated with chronic thermal stress. Transcripts involved in heat stress and regulatory responses (i.e., *hsp90ab1*, *hspa8*, *gpx1*, *nkcc1a* and *nupr1*) exhibited differential abundance at 20°C and higher suggesting a thermal threshold at the transcript level. Elevated levels of plasma cortisol and glucose after the exhaustive exercise and air exposure stressor, suggests increased activation of the HPI axis at 20°C and higher. Increases in muscle lactate also suggests increased anaerobic metabolic processes at 20°C and higher. Additionally, standard metabolic rate was higher at elevated temperatures (20°C), and fish experienced extended recovery times at high acclimation temperatures (15 and 20°C), indicating a more pronounced response to the acute stressor at elevated temperatures. Overall, the ability for juvenile brook trout to acclimate for 3 weeks to temperatures beyond 20°C appears to be reduced.

### Cellular-level response to chronic temperature exposure

Juvenile brook trout at temperatures above 20°C showed a cellular heat shock response in gill and liver tissues. Heat shock proteins are part of a ‘classic’ temperature-induced cellular stress response and play a critical role in reducing and repairing damage to cellular proteins that arises from physical or chemical stress (Iwama et al., 1998; Somero, 2010; Currie, 2011). When temperatures begin to approach a species’ thermal limit, heat shock protein expression is often increased (Currie, 2011). In the present study, both *hsp90ab1* and *hspa8* (*hsc70*) mRNA levels were significantly elevated at 23 and 25°C compared to temperatures ≤15°C. Similarly, in the gill tissue there was a significant increase in abundance of *hsp90ab1* mRNA at 25°C compared to temperatures ≤15°C. Other salmonid species such as, arctic char (*Salvelinus alpinus*), sockeye salmon (*Oncorhynchus nerka*), and pink salmon (*O. gorbuscha*), have had elevated mRNA levels of *hsp90ab1* and *hspa8* at temperatures ≥19°C (Quinn et al., 2011; Jeffries et al., 2012a, 2014; Akbarzadeh et al., 2018). As both *hsp90ab1* and *hspa8* are constitutive isoforms (Iwama et al., 1998), changes in their expression would be expected in chronic events such as thermal acclimation, and our results further suggest that brook trout are activating a chronic cellular stress response above 20°C as evidenced by increased mRNA abundance at these temperatures.

As temperature surpassed 20°C, an apparent thermal threshold was reached for genes involved in the oxidative stress response in gill and liver tissues. Glutathione peroxidase 1 is responsible for catalyzing the reduction of H_2_O_2_ into H_2_O or alcohol and thereby prevents damage caused by reactive oxygen species (ROS; Sattin et al., 2015). As metabolic rate increases with temperature, there is an increase in oxidative phosphorylation in the mitochondria, potentially resulting in elevated production of ROS (Davidson and Shiestl, 2001). The SMR of brook trout increased significantly with temperature in the present study, potentially resulting in elevated ROS production, and thus may be responsible for the increase in abundance of *gpx1* mRNA observed in the gill at temperatures above 20°C. In contrast, a significant increase in the abundance of *gpx1* mRNA in the liver was observed at cooler temperatures (i.e., 5 and 10°C) compared to fish held at 20°C. Multiple Antarctic fishes (e.g., *Champsocephalus gunnari, Chaenocephalus aceratus, Pseudchaenichthys georgianus, Dissostichus eleginoides,* and *Notothenia* rossi) have exhibited increased glutathione peroxidase activity in the heart and liver tissues compared to the gills and muscle tissues (Ansaldo et al. 2000). An increase of glutathione peroxidase in the liver may be due to the dependence of aerobic metabolism on the oxidation of unsaturated fats in the liver, fats that are susceptible to oxygen radical attack (Roberfroid and Calderon, 1995). Previous work on this same group of brook trout found that hepatosomatic index (comparison of liver mass to body mass) was lowest at 23 and 25°C and higher at cooler temperatures (5°C), indicating a higher fat content at cooler temperatures (Morrison et al., 2020). The relationship between higher fat content in the liver and higher ROS attack is supported by results from Morrison et al. (2020) and may explain why liver tissue exhibited increased abundance of *gpx1* mRNA in our study. Overall, *gpx1* mRNA abundance differed across gill and liver tissue where increased abundance was observed at warmer temperatures for the gill tissue in contrast to increased abundance at cooler temperatures for the liver tissue.

The mRNA abundance of genes involved in cellular processes, such as cell growth, and metabolic processes (e.g., glycogen production), exhibited peak abundance levels at 20°C, levels that subsequently decreased at higher temperatures. Because some cellular responses can exhibit a peak near or prior to detrimental physiological changes, the peak mRNA transcript levels at 20°C may suggest that it is near a sub-lethal threshold. Nuclear protein 1 is involved in the regulation of cell growth and apoptosis (Mallo et al., 1997) and plays a role as a transcription factor (Momoda et al., 2007). The *nupr1* mRNA abundance in the liver was highest in juvenile brook trout held at 20°C, and then decreased by 2-fold in fish held at 25°C in the present study. This decline in *nupr1* mRNA abundance in fish held at 25°C compared to 15 and 20°C suggests a potential sub-lethal threshold for *nupr1* at temperatures above 20°C. Additionally, higher mortality rates were observed in fish held at 25°C, further supporting the shutdown of certain cellular processes at temperatures beyond 23°C. This increased abundance of mRNA at 20°C was also observed for liver *g6pc*, which plays a role in glycogen metabolism, where levels peaked at 20°C. Therefore, elevation of *nupr1* and *g6pc* abundance at 20°C and decreased abundance at 25°C is consistent with a shift in the cellular processes being activated at the highest acclimation treatments.

### Whole-animal responses to chronic temperature exposure

The acclimation temperature affected the ability of the juvenile brook trout to mount a stress response to an acute stressor. Plasma cortisol levels typically increase in fish between 30–60 min following exposure to an acute stressor (Biron and Benfey, 1994; Benfey and Biron, 2000), which can lead to gluconeogenesis, thereby elevating levels of plasma glucose (Wendelaar Bonga, 1997). In our study, the cortisol response to an acute stressor was impaired or delayed in fish exposed to lower temperatures (5 and 10°C), as there was no apparent increase in plasma cortisol levels 30 min following stressor exposure (acute-stress group) or after 24 h of recovery (acute-recovery group). Similarly, there was no change in plasma glucose levels in response to the acute stressor in fish held at 5 and 10°C. In other studies, peak plasma cortisol levels for fish exposed to lower temperatures were also significantly delayed, possibly due to reduced enzymatic activity at cold temperatures (Van Ham et al., 2003; Louison et al., 2017). Conversely, plasma cortisol levels were significantly elevated 30 min post-exposure to the exhaustive exercise and air exposure stressor in fish held at temperatures ≥15°C. Notably, plasma cortisol levels did not return to pre-stress levels (i.e., unhandled group) 24 h post-stressor exposure (20 and 23°C), suggesting that the fish had not fully recovered from the stressor at the higher temperatures. Plasma glucose levels showed similar increases post-stressor exposure in fish held at temperatures ≥15°C, with the exception that glucose levels returned to pre-stress levels for fish acclimated to 20 and 23°C. This pattern of significantly elevated circulating cortisol levels at higher temperatures post-exposure to an exhaustive exercise stressor has been displayed in several other studies (e.g., Jain and Farrell, 2003; Suski et al., 2003; Meka and McCormick, 2005; Suski et al., 2006; McLean et al., 2016). For example, rainbow trout (*Oncorhynchus mykiss*) in southwest Alaska that were angled during a warmer year compared with a cooler year (13.2°C vs. 9.8°C, respectively) exhibited significantly increased plasma cortisol concentrations post-angling event (Meka and McCormick, 2005). Our data suggests that at higher temperatures, especially post-stressor exposure, there was an elevated stress response in brook trout demonstrated by increased plasma cortisol and glucose. However, possibly due to enzymatic properties, it is unclear if there was sufficient time for plasma cortisol and glucose to reach peak values at the cooler temperatures (5 and 10°C).

Lactate is a by-product of anaerobic metabolism and increased concentrations of lactate in the white muscle of fish can be caused by extensive exercise and activity (Wood et al., 1983). Across all acclimation temperatures, the exhaustive exercise and air exposure stressor led to a significant transient increase in lactate in the white muscle 30 min post-stressor exposure, that returned to or below pre-stress (i.e., unhandled) levels 24 h later. Several studies on a range of fishes, including brook trout, have demonstrated increased muscle lactate or plasma lactate concentrations when fish were exposed to exercise and air exposure (Beggs et al., 1980; Ferguson and Tufts, 1992; Booth et al., 1995; Milligan, 1996; Farrell et al., 2001; Kieffer et al., 2011; Landsman et al., 2011). Interestingly, our results showed a decrease in muscle lactate below pre-stress levels after 24 h recovery in fish held at 23°C group. A decrease in muscle lactate can be caused by a release of lactate into the blood or because the lactate in the muscle is recycled *in situ* for glycogenesis (Milligan and Wood, 1986; Milligan and Girard, 1993; Kieffer et al., 1994; Milligan, 1996). Therefore, the observed decrease in muscle lactate after 24 h recovery in the 23°C group may be a result of its release into the blood stream or recycling through glycogenesis to help the fish return its energy stores to pre-stress levels. Our results suggest that the exhaustive exercise and air exposure induced anaerobic respiration as exhibited by the increase in muscle lactate across all temperature groups.

We found some evidence that osmoregulation in brook trout was impacted at elevated temperatures as shown by increased levels of plasma osmolality and changes in mRNA abundance of *nkcc1a*. Plasma osmolality can estimate the osmoregulatory ability of fishes as an indicator of ion balance, particularly for circulating concentrations of Na^+^ and Cl^−^ (McDonald and Milligan, 1997). For the brook trout that were exposed to the exhaustive exercise and air exposure stressor, we observed a significant transient increase in plasma osmolality 30 min post-stressor exposure that returned to pre-stress levels (i.e., unhandled) in fish held at 15 and 20°C. After exposure to an acute stressor, increased cardiac output would lead to increased blood perfusion at the gill (Mazeaud and Mazeaud, 1981; Sopinka et al., 2016). However, there is a trade-off between improving oxygen uptake at the expense of increasing permeability to water and ions, termed the osmorespiratory compromise (Randall et al., 1972; Nilsson, 2007; Onukwufor and Wood, 2018). This compromise may aid in explaining why there was no increase in osmolality in the acute-stress group at 23°C but there was a decline 24 h later. At a molecular level, *nkcc1a,* a gene involved in ion regulation showed increased mRNA abundance in the gill at low temperatures, with decreased abundance at higher temperatures. Because the Na^+^-K-Cl^−^ cotransporter is involved in active ion absorption or secretion across cellular membranes in gills (Hiroi et al., 2008), this may further suggest that brook trout osmoregulatory ability was potentially impacted by increased temperatures.

Both SMR and MMR changed with acclimation temperature, but AS was not significantly affected in juvenile brook trout. The SMR and MMR increased with temperature, with the highest levels evident in fish held at 23°C. There is a strong positive relationship of water temperature on metabolic rate in ectotherms (Fry, 1971; Hulbert and Else, 2004; Brett, 1964; Beamish, 1978), therefore increased SMR at higher temperatures was expected in brook trout. Likely due to the nearly parallel increase in SMR and MMR, AS was not significantly affected by acclimation temperature. Stable AS values were also observed in Nile perch (*Lates niloticus*) that were acclimated for three weeks at 27, 29, and 31°C (Nyboer and Chapman, 2017). A similar trend of stable AS across temperatures has also been observed in Chinook salmon (*O. tshawytscha*, Poletto et al., 2017) and pink salmon (*O. gorbuscha*, Clark et al., 2011). The maintenance of similar AS in fish acclimated to different temperatures may be evidence for metabolic compensation (Donelson and Munday, 2012). If the energy available to allocate to other processes (AS) remains constant, the body may have to adjust and metabolically compensate to maintain energy, oxygen, heart rate, and other vital processes (Eliason and Farrell, 2016). Overall, our metabolic data reflects an increase in metabolic activity at higher temperatures with possible metabolic compensation as indicated by constant AS across all temperatures.

Time to recovery was highest at 15 and 20°C, while 23°C displayed the lowest recovery time of all groups. The fish acclimated to 23°C exhibited the shortest recovery time, despite having the highest SMR and evidence of cellular impairment. This short recovery time at 23°C does not seem to be due to this group having a higher SMR, as the AS between all groups across the study was not significantly different. Considering the constant AS across groups it is possible that this quick recovery time in the 23°C may be due to respiratory alkalosis and metabolic suppression. Metabolic rate depression has been a strategy used by other animals to combat adverse environmental conditions (Hochachka and Guppy, 1987). For example, goldfish (*Carassius auratus*) can suppress their metabolism up to 30% of the aerobic metabolic rate at elevated environmental temperatures (Van Waversveld et al., 1988). Post exhaustive exercise and air exposure, may lead to elevated blood CO2 levels and lower blood pH (Wang et al., 1994; Milligan, 1996) and this can be exacerbated at high temperatures. Metabolic suppression may also be used to limit further carbon dioxide build up in the blood (acidosis) to regulate the blood pH of fishes (Claiborne, 1998). Therefore, it is possible that the short recovery time in the fish from the 23°C acclimation group could be attributed to metabolic suppression.

### Conclusion

We demonstrated that the acclimation ability of brook trout to temperatures ≥20°C is impaired as shown by changes in mRNA transcript abundance, standard metabolic rate, and responses to exhaustive exercise and air exposure. These findings are consistent with previous work that suggested that upper temperatures limiting habitat use in brook trout is around 21– 23.5°C (reviewed in Smith and Ridgway 2019). Our study indicated that there is a sub-lethal threshold between 20–23°C that shows when chronic temperatures may begin to adversely impact the physiological performance of brook trout. Previous work on this same group of brook trout also showed a sub-lethal threshold between 20–23°C as fish were no longer able to increase their critical thermal maxima with acclimation temperature and there was increased plasma lactate levels indicating anaerobic metabolism (Morrison et al. 2020). Furthermore, Chadwick and McCormick (2017) demonstrated that brook trout growth is limited by higher temperatures especially those above 23°C, and that may play a role in driving their distribution. Collectively, these studies in combination with the present study suggest that there is a limitation to the ability of brook trout to cope with chronic temperatures ≥20°C, which can potentially provide a benchmark for understanding the ability of some brook trout populations to persist in the wild in the future.

## Acknowledgments

We thank Andrew Chapelsky, Scott Morrison, and Jamie Card for assisting with the experiments and Kerry Wautier for assistance in rearing of the fish. Thank you to Dr. Gary Anderson for the use of his laboratory for osmolality analyses and Alex Schoen for help with the glucose assay.

