## Supplemental Data for "Molecular and physiological responses predict acclimation limits in juvenile brook trout (*Salvelinus fontinalis*)"

**Supplementary Information**

**Table S1.** Transcript information for the thirteen target and three reference genes retrieved from a previously published brook trout transcriptome in Sutherland et al., (2019).

| Subject | Length (bp) | Species | Database | Accession no. | Score | E. value | % Identity |
|---|---|---|---|---|---|---|---|
| QSF_HSP70.6.8 | 1932 | *Oncorhynchus mykiss* | Swissprot | HSP70_ONCMY | 2732 | 0 | 86.5 |
| QSF_SERPH.3.3 | 1233 | *Salmo salar* | Refseq_rna | NM_001139968.1 | 840 | 0 | 92.6 |
| QSF_ATP1A3.7.8 | 972 | *Oncorhynchus mykiss* | Refseq_rna | NM_001124630.1 | 458 | 0 | 84.8 |
| QSF_G6PC.1.2 | 1059 | *Felis catus* | Swissprot | G6PC_FELCA | 1059 | $1e^{-25}$ | 49.9 |
| QSF_CIRBP.4.24 | 468 | *Salmo salar* | Refseq_rna | NM_001139676.1 | 152 | $7e^{-79}$ | 88 |
| QSF_GPX1.1.4 | 255 | *Oncorhynchus mykiss* | Refseq_rna | NM_001124525.1 | 418 | 0 | 91.2 |
| QSF_HSP90B.2.6 | 1161 | *Salmo salar* | Refseq_rna | NM_001123532.1 | 818 | 0 | 95.5 |
| QSF_LOC10117185.1.1 | 228 | *Oryzias latipes* | Refseq_protein | XP_004086462.1 | 312 | $1e^{-31}$ | 76.4 |
| QSF_RL7.4.13 | 942 | *Salmo salar* | Refseq_rna | NM_001140480.1 | 523 | 0 | 93.6 |
| QSF_RS9.1.4 | 267 | *Rattus norvegicus* | Swissprot | RS9_RAT | 200 | $1e^{-16}$ | 97.4 |
| QSF_RL8.1.6 | 774 | *Danio rerio* | Swissprot | RL8_DANRE | 1267 | $1e^{-177}$ | 93.8 |
| QSF_IRK8.1.1 | 1266 | *Salmo salar* | Refseq_rna | NM_001140360.1 | 1828 | 0 | 97.2 |
| QSF_NKCC1A.1.2 | 3453 | *Salmo salar* | Refseq_rna | NM_001123683.1 | 479 | 0 | 94.1 |
| QSF_LOC100136366.1.1 | 4557 | *Salmo salar* | Refseq_rna | NM_001123534.1 | 4750 | 0 | 96.9 |
| QSF_LOC100136607.1.1 | 561 | *Oncorhynchus mykiss* | Refseq_rna | NM_001124597.1 | 633 | 0 | 97.6 |
| QSF_LOC101477634.1.1 | 681 | *Maylandia zebra* | Refseq_rna | XM_004548350.1 | 259 | $1e^{-148}$ | 84.8 |

**Table S2.** Results of one-way analysis of variance (ANOVA) and Kruskal-Wallis test for the mRNA abundance of genes measured in gill and liver tissue of juvenile brook trout (*Salvelinus fontinalis*) acclimated to 5, 10, 15, 20, 23, and 25˚C.

| Tissue | Gene | Df | Sum of Squares | Mean of Squares | *F* values/ Chi-squared | *P*-value |
|---|---|---|---|---|---|---|
| Gill | *atp1a3* | 5, 59 | 24.39 | 4.878 | 3.674 | **0.006** |
| | *irk8* | 5, 59 | 14.60 | 2.920 | 1.739 | 0.142 |
| | *nkcc1a* | 5, 59 | 35.33 | 7.066 | 5.069 | < **0.001** |
| | *vatb* | 5, 59 | - | - | 2.268 | 0.811 |
| | *vate1* | 5, 59 | 10.28 | 2.056 | 1.475 | 0.213 |
| | *cftr* | 5, 59 | 2.96 | 0.591 | 0.4 | 0.846 |
| | *cirbp* | 5, 59 | 14.83 | 2.966 | 1.469 | 0.216 |
| | *gpx1* | 5, 59 | 6007 | 1201.4 | 5.735 | < **0.001** |
| | *serpinh1* | 5, 59 | 10.55 | 2.111 | 1.933 | 0.104 |
| | *hspa8* | 5, 59 | 11.94 | 2.387 | 2.252 | 0.063 |
| | *hsp90ab1* | 5, 59 | 25.33 | 5.066 | 4.321 | **0.002** |
| Liver | *g6pc* | 5, 59 | 67.56 | 13.51 | 3.123 | **0.016** |
| | *gpx1* | 5, 59 | 36.22 | 7.245 | 4.924 | **0.001** |
| | *hspa8* | 5, 59 | 36.59 | 7.318 | 8.828 | < **0.001** |
| | *hsp90ab1* | 5, 59 | 54.31 | 10.86 | 15.14 | < **0.001** |
| | *nupr1* | 5, 59 | 4444 | 888.8 | 4.918 | **0.001** |
| | *serpinh1* | 5, 59 | 4.63 | 0.9258 | 0.573 | 0.720 |

*atp1a3, $Na^+/K^+$-transporting ATPase subunit alpha-3*; *irk8, ATP-sensitive inward rectifier $K^+$ channel 8*; *nkcc1a, $Na^+/K^+/2Cl^-$ co-transporter-1-a*; *vatb, B subunit of V-type H-ATPase*; *vate1, E 1 subunits of V-type H-ATPase*; *cftr, cystic fibrosis transmembrane conductance regulator*; *cirbp, cold-inducible RNA-binding protein*; *gpx1, glutathione peroxidase-like peroxiredoxin*;

*serpinh1, serpinh1*; *hspa8, heat shock cognate 71 kDA protein*; *hsp90ab1, heat shock protein 90-beta*; *g6pc, glucose-6-phosphatase*; *nupr1, nuclear protein-1*.

Data were analyzed using a one-way ANOVA except if data were not equally variable or normally distributed, in which case a Kruskal-Wallis test was used; significant *P*-values are represented as bolded text

**Table S3.** Results of the two-way analysis of variance (ANOVA) for muscle lactate, as well as plasma cortisol and glucose of juvenile brook trout (*Salvelinus fontinalis*) acclimated to five temperatures (5, 10, 15, 20, and 23˚C) and exposed to one of three treatments (unhandled, acute stress, and acute-stress recovery).

| **Variable** | **Group** | **Df** | **Sum of Squares** | **Mean of Squares** | ***F* value** | ***P*-value** |
|---|---|---|---|---|---|---|
| Muscle lactate | Treatment | 2 | 13.135 | 6.568 | 125.415 | < **0.001** |
| | Temperature | 4 | 4.107 | 1.027 | 19.606 | < **0.001** |
| | Treatment × Temperature | 8 | 4.106 | 0.513 | 0.513 | < **0.001** |
| Plasma cortisol | Treatment | 2 | 37274 | 18637 | 46.15 | < **0.001** |
| | Temperature | 4 | 6593 | 1648 | 4.08 | **0.004** |
| | Treatment × Temperature | 8 | 13919 | 1740 | 4.31 | < **0.001** |
| Plasma glucose | Treatment | 2 | 2.9573 | 1.4787 | 64.803 | < **0.001** |
| | Temperature | 4 | 0.7413 | 0.1853 | 8.122 | < **0.001** |
| | Treatment × Temperature | 8 | 1.2648 | 0.1581 | 6.929 | < **0.001** |

Significant *P*-values are represented as bolded text

**Table S4.** Results of the one-way ANOVA and Kruskal-Wallis test for standard metabolic rate (SMR) and maximum MR (MMR), recovery time, and aerobic scope (AS) of juvenile brook trout (*Salvelinus fontinalis*) acclimated to 5, 10, 15, 20, and 23˚C.

| **Variable** | **Df** | **Sum of Squares** | **Mean of Squares** | ***F* value/ Chi-Squared** | ***P*-value** |
|---|---|---|---|---|---|
| **SMR** | 4, 40 | - | - | 30.43 | < **0.001** |
| **MMR** | 4, 40 | 1.272 | 0.318 | 8.84 | < **0.001** |
| **Recovery Time** | 4, 40 | 506.7 | 126.7 | 27.69 | < **0.001** |
| **AS** | 4, 40 | 0.598 | 0.149 | 1.29 | 0.292 |

Significant *P*-values are represented as bolded text

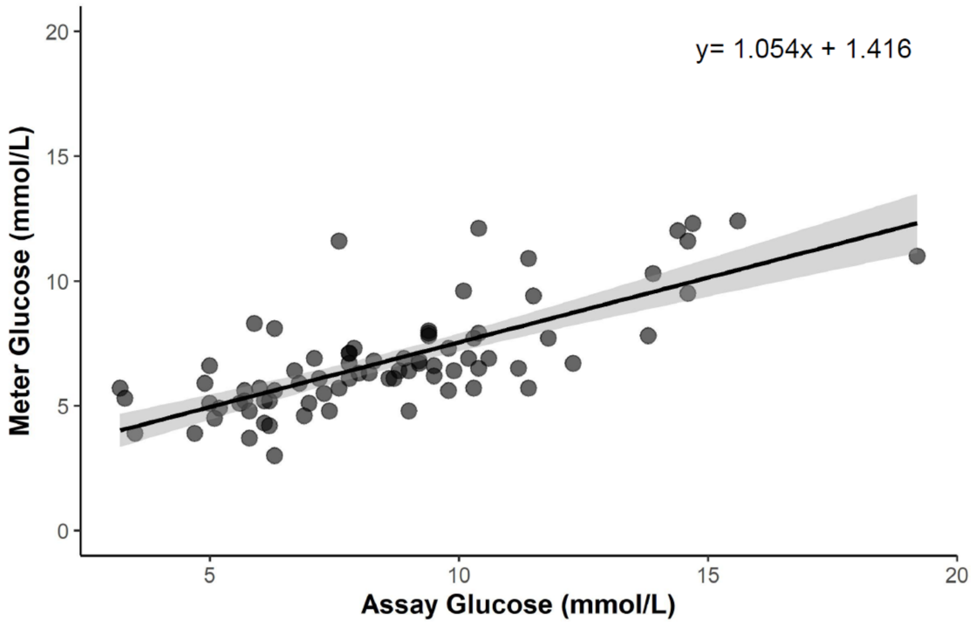


**Figure S1.** Comparison of glucose values from the hexokinase kinetic assay and the glucose meter using a linear regression (y = 1.054x + 1.416) for juvenile brook trout (*Salvelinus fontinalis*) held at temperatures spanning their thermal distribution (n = 8–10 per temperature treatment).

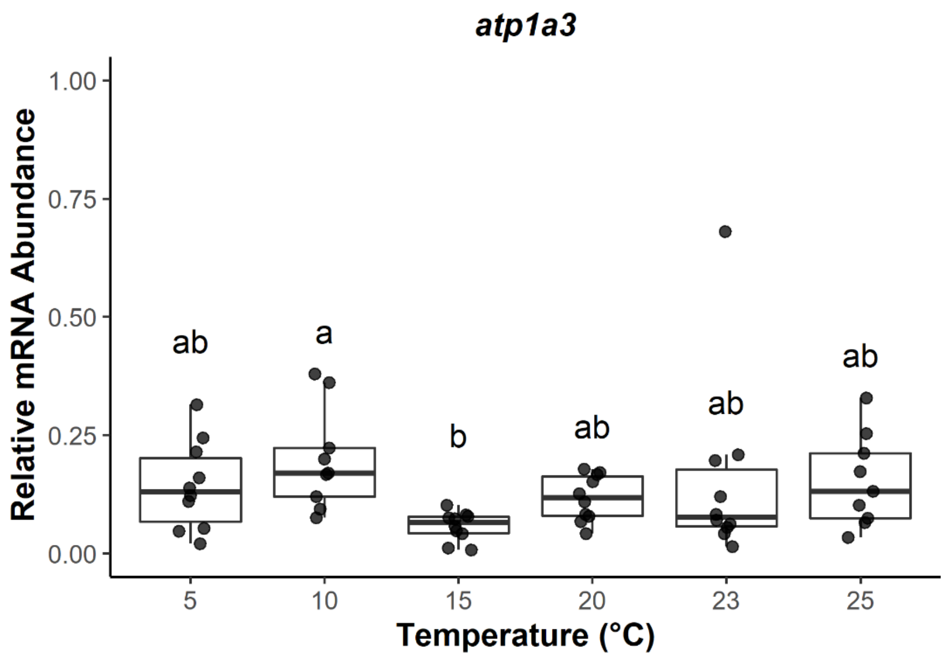


**Figure S2.** Transcript abundance of *atp1a3* (*$Na^+$/$K^+$-transporting ATPase subunit alpha-3*) in gill tissue for juvenile brook trout (*Salvelinus fontinalis*) held at temperatures spanning their thermal distribution (*n* = 8–10). Fish were held for 21 days at the respective acclimation temperature, with the exception of fish held at 25˚C, where fish were sampled after 11 days (see text for details). Groups that do not share a letter are significantly different from one another (one-way ANOVA, $P < 0.05$; see Table S2). Horizontal bars in the boxplot represent the median response value and the 75 and 25% quartiles. Whiskers represent ± 1.5 times the interquartile range, and each dot represents and individual response value.

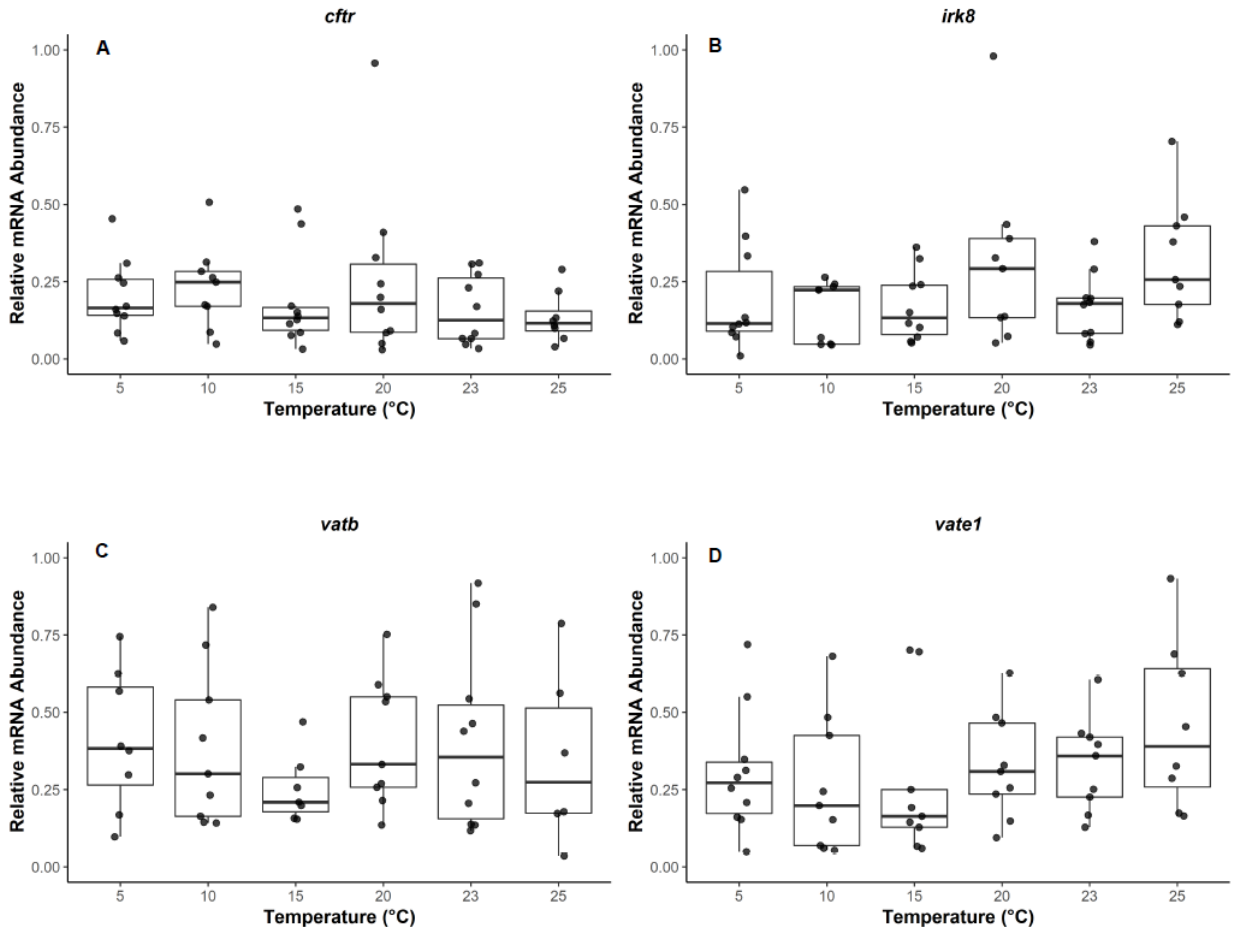


**Figure S3.** Transcript abundance ion regulation biomarkers in gill tissue for juvenile brook trout (*Salvelinus fontinalis*) held at temperatures spanning their thermal distribution *(n* = 8–10*)*. Fish were held for 21 days at the respective acclimation temperature, with the exception of fish held at 25˚C, where fish were sampled after 11 days (see text for details). Groups that do not share a letter are significantly different from one another (one-way ANOVA, $P < 0.05$; see Table S2). Horizontal bars in the boxplot represent the median response value and the 75 and 25% quartiles. Whiskers represent ± 1.5 times the interquartile range, and each dot represents and individual response value. *cftr, cystic fibrosis transmembrane conductance regulator*; *irk8, ATP-sensitive inward rectifier* $K^+$ *channel 8*; *vatb and vate1, B and E1 subunits of V-type H-ATPase*.

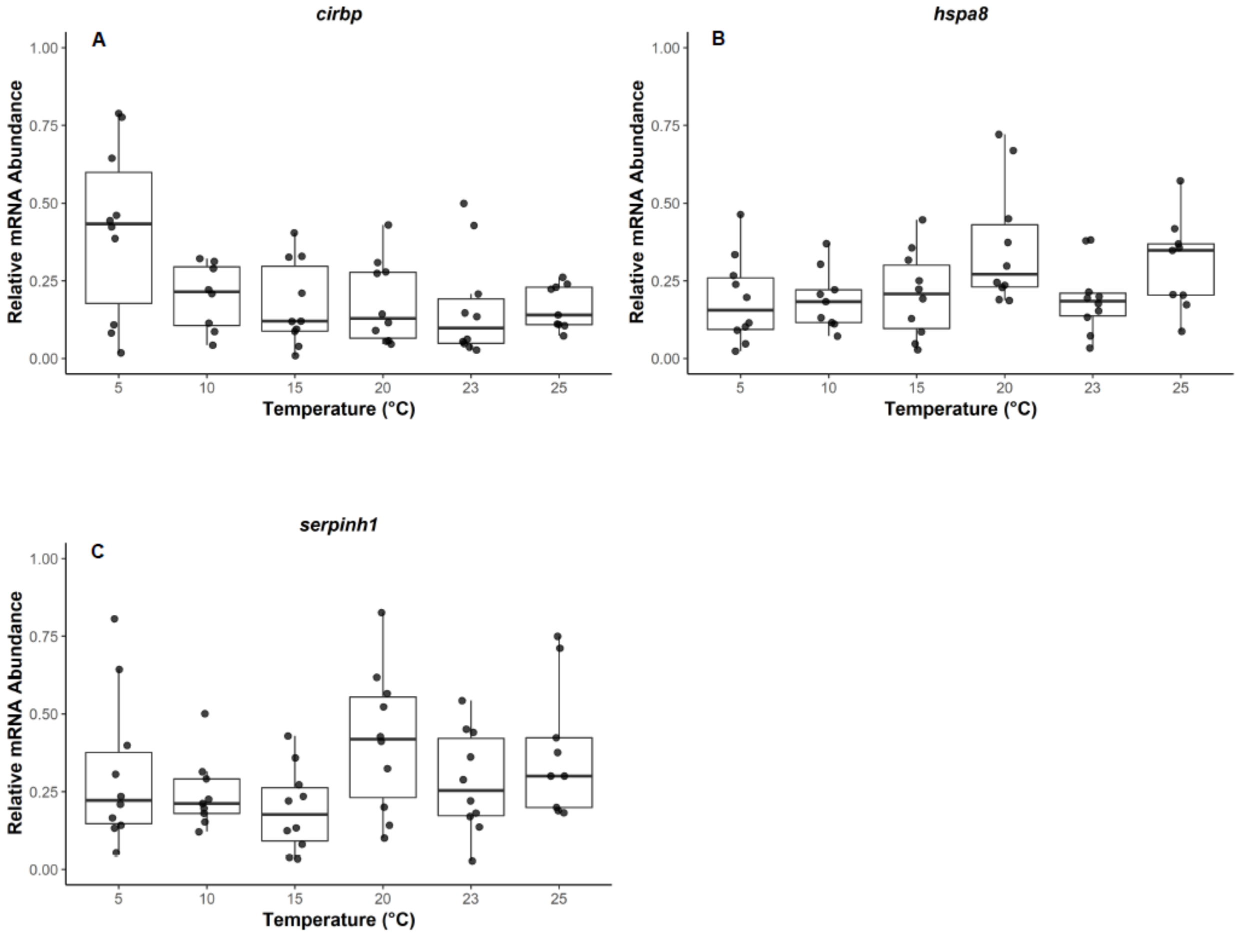


**Figure S4.** Transcript abundance of thermal stress biomarkers in gill tissue for juvenile brook trout (*Salvelinus fontinalis*) held at temperatures spanning their thermal distribution (*n* = 8–10). Fish were held for 21 days at the respective acclimation temperature, with the exception of fish held at 25˚C, where fish were sampled after 11 days (see text for details). Groups that do not share a letter are significantly different from one another (one-way ANOVA, $P < 0.05$; see Table S2). Horizontal bars in the boxplot represent the median response value and the 75 and 25% quartiles. Whiskers represent ± 1.5 times the interquartile range, and each dot represents and individual response value. *cirbp*, *cold-inducible RNA-binding protein*; *hspa8*, *heat shock cognate 71kDA protein*; *serpinh1*, *serpinh1*.

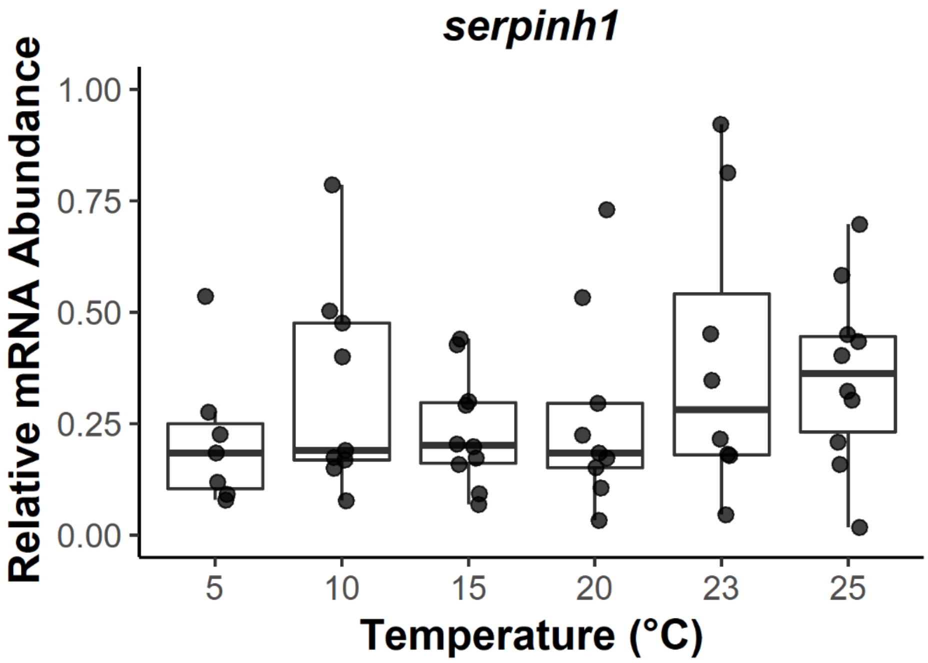


**Figure S5.** Transcript abundance of *serpinh1* (*serpinh1*) in liver tissue for juvenile brook trout (*Salvelinus fontinalis*) held at temperatures spanning their thermal distribution ($n$ = 8–10). Fish were held for 21 days at the respective acclimation temperature, with the exception of fish held at 25˚C, where fish were sampled after 11 days (see text for details). Groups that do not share a letter are significantly different from one another (one-way ANOVA, $P < 0.05$; see Table S2). Horizontal bars in the boxplot represent the median response value and the 75 and 25% quartiles. Whiskers represent ± 1.5 times the interquartile range, and each dot represents and individual response value.
